# Programming N-linked glycan composition through de novo protein design

**DOI:** 10.64898/2026.10.07.757460

**Authors:** Margaret C. Lunn-Halbert, Joel D. Allen, Rohith Krishna, Anna Manchenko, Xinting Li, Edward Lopatto, Jerome Pinkner, Carolina Moller, Woody Ahern, Johnny Mendoza, Alex Kang, Hannah Nguyen, Emily Joyce, Asim Bera, Scott J. Hultgren, Evgeni Sokurenko, Max Crispin, David Baker, Neil P. King

## Abstract

N-linked glycans are ubiquitous post-translational modifications that mediate diverse biological processes, including cell signaling and adhesion, molecular recognition, and host-pathogen interactions. Despite their importance, N-linked glycans have largely remained beyond the reach of biomolecular design due to a lack of methods capable of jointly modeling glycans and protein structure. Here, we develop a general all-atom computational pipeline for creating de novo glycoproteins in which the amino acid sequence encodes both the location and composition of N-linked glycans. The method uses three-dimensional protein structure to tune glycan accessibility to processing enzymes in the eukaryotic secretory pathway. Systematic increases in steric restriction progressively reduced processing, shifting glycans from heterogeneous complex-type structures toward increasingly underprocessed oligomannose-type species resembling those found on many viral glycoproteins. Despite this programmed restriction, the resulting oligomannose glycans remained accessible for biological recognition, inhibiting uropathogenic *Escherichia coli* adhesion to human bladder cells and engaging mannose-binding lectin. These results establish steric control of N-glycan processing as a genetically encodable design principle and extend computational protein design to functional glycoproteins with programmed composition.

## INTRODUCTION

More than half of mammalian proteins are estimated to be glycosylated, underscoring the central role of glycans in biology^1^. Glycans mediate protein folding and trafficking, cell signaling and adhesion, molecular recognition, and immune discrimination between self and non-self^2^. Mammalian glycans range from simple monosaccharide modifications to large glycosaminoglycans, and differences in glycan structure can profoundly alter glycoprotein function. Consequently, considerable effort has been directed toward engineering glycan composition for biotechnological applications. For example, modifying the Fc N-glycan in immunoglobulins can modulate antibody effector functions such as Fc receptor binding and complement activation^3–7^. Similarly, oligomannose-type glycans abundant on yeast and immaturely processed glycoproteins are recognized by lectins of the innate immune system and can enhance immune responses when incorporated into vaccines^8^. Nevertheless, current approaches to glycan engineering, including glycoengineered cell lines and small-molecule inhibitors of glycan-processing enzymes, remain fundamentally limited. These methods generally act globally rather than at individual glycosylation sites, making it difficult to control the composition of a specific glycan without perturbing glycosylation throughout the entire cell. For example, kifunensine can enrich oligomannose-type N-glycans by inhibiting ER mannosidase I, but it non-specifically affects all susceptible glycans and is poorly suited to commercial-scale biologics manufacturing^9,10^.

N-linked glycan synthesis begins co-translationally, when oligosaccharyltransferase (OST) transfers a preassembled dolichol pyrophosphate-linked Glc_3_Man_9_GlcNAc_2_ (Glc3Man9) oligosaccharide onto the asparagine of an N-X-S/T sequon (X ≠ P) in proteins destined for secretion or membrane insertion^11–14^. Following protein folding, glucose and mannose residues are sequentially removed by glycosidases and mannosidases, generating a common Man_5_GlcNAc_2_ (Man5) core that serves as the substrate for further processing. In most mature mammalian glycoproteins, N-glycans are processed beyond this stage through additional trimming and subsequent extension reactions that incorporate monosaccharides such as N-acetylglucosamine, fucose, galactose, and sialic acid^15^. This late-stage processing is not directly templated by the amino acid sequence, and the final glycan composition is influenced by factors including species, cell type, glycosyltransferase expression, enzyme localization, and precursor availability^16,17^. As a result, mature complex-type N-glycans often exhibit substantial microheterogeneity in structure and composition.

The local structural context surrounding an N-linked glycan can also influence processing. This principle is evident in many viral glycoproteins, where dense glycan shields sterically hinder processing enzymes from accessing their substrates^16,18–22^. The HIV-1 envelope glycoprotein (Env) is a prominent example: crowding among neighboring glycans creates an “intrinsic mannose patch” in which processing is arrested before conversion to complex-type structures, resulting in the persistence of underprocessed oligomannose-type glycans on mature Env^23^. Because oligomannose-type glycans are common on pathogens but uncommon on mature host glycoproteins, the mammalian immune system has evolved soluble and cell-associated lectins that recognize these structures as pathogen-associated molecular patterns^8,24–26^. Mannose-binding lectin, for example, recognizes mannose-rich glycan arrays and can initiate complement activation through the lectin pathway. Conversely, some microbes use lectin-like adhesins to exploit host glycans during colonization. Uropathogenic *E. coli*, for example, uses the fimbrial tip adhesin FimH to bind mannosylated receptors on uroepithelial cells, a process countered by the abundant urinary protein uromodulin, which presents genetically encoded oligomannose glycans as receptor decoys^27–29^. These observations identify steric restriction as a mechanism for controlling N-glycan composition and suggest that programmable access to oligomannose-type glycans could enable the design of new glycoproteins with glycan-driven biological activity.

While computational protein design is an established route to atom-level control over de novo protein structures, advances in “all-atom” modeling have only recently begun to extend this design paradigm beyond proteins alone. Methods such as RoseTTAFold diffusion All-Atom (RFdiffusionAA), LigandMPNN, and AlphaFold3 (AF3) now enable structure prediction and design for systems containing nucleic acids, small molecules, and covalent modifications including glycans^30–33^. However, the use of such methods to design de novo glycoproteins has not yet been demonstrated.

Here we develop a general computational approach for designing de novo N-linked glycoproteins with programmable glycan location and composition. We show that the three-dimensional structures of de novo glycoproteins can genetically encode oligomannose-type glycans through programmed steric restriction, and that the resulting glycoproteins function as bacterial adhesion inhibitors or immune lectin engagers.

## RESULTS

### Computational design, expression, and characterization of de novo glycoproteins

To design de novo glycoproteins with a potential N-linked glycosylation site (PNGS), we used recently described deep-learning methods capable of modeling non-protein atoms^32^. Beginning with RFdiffusionAA^32^, we generated structurally diverse protein backbones around an explicitly modeled asparagine-linked Man9 glycan^34^ (**Figure 1a**). We then assigned amino acid sequences to these backbones using the glycan-aware LigandMPNN (**Figure 1b**) and evaluated the resulting designs using AlphaFold2 (AF2) without an explicitly modeled glycan^31,35^. Because NXT sequons are associated with higher glycan occupancy than NXS sequons, we ensured that most +2 positions were threonine^19,36,37^. 257 designs predicted with high confidence and low deviation from the diffused backbone [predicted aligned error (pAE) < 10 and predicted local distance difference test (pLDDT) > 80; RMSD <3 Å] were selected for experimental characterization (**Figure 1c**; LC-MS/MS detected amino acid sequences provided in **Supplementary Data S1**).

**Fig. 1:**
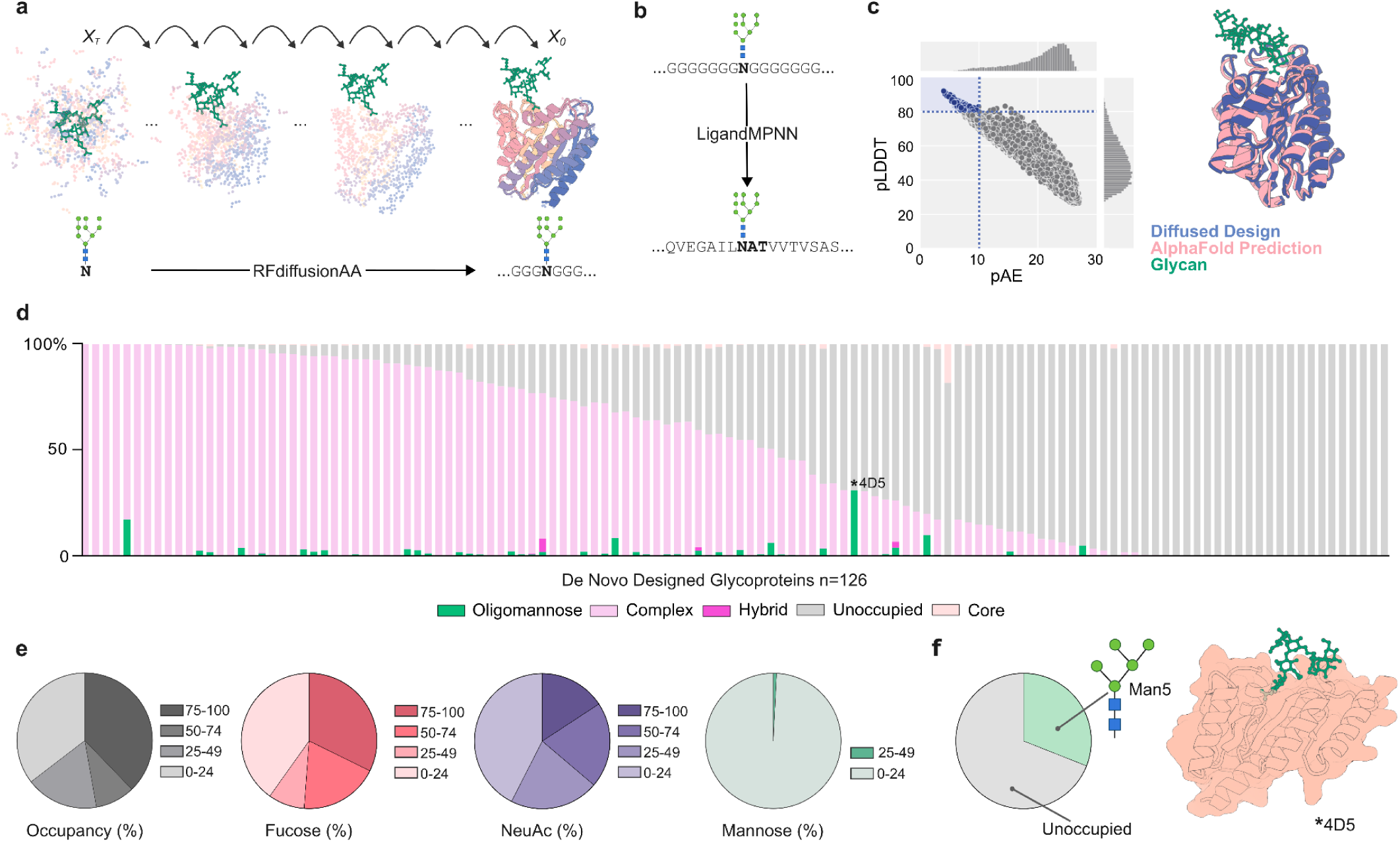
Computational design, expression, and characterization of de novo glycoproteins. a, Protein backbones were generated using RFdiffusionAA^32^. Input structures contained a modeled Man9 glycan attached to a single asparagine residue or full PNGS motif. Input glycans were modeled using GlycoShape^34^. **b,** Protein backbones were used as inputs to LigandMPNN to assign sequences containing a PNGS motif utilizing an explicitly modeled Man9 glycan^31^. **c,** Sequences were predicted using AF2 and filtered on pLDDT and pAE metrics. **d,** LC-MS/MS analysis of de novo glycoproteins. Each bar represents a unique glycoprotein. Glycans were binned by complex (light pink), hybrid (magenta), oligomannose (green), or unoccupied (gray). The oligomannose glycoprotein highlighted in panel d is marked with an asterisk (*). **e,** LC-MS/MS glycosylation analysis of overall occupancy (gray), fucosylation (red), sialylation (purple), or oligomannose-type glycosylation (green) by percentage, averaged across all individual glycoproteins (n=126). **f,** *Left,* Site-specific glycan analysis of a single oligomannose-type glycoprotein with occupancy >25%. Total glycosylation proportion of unoccupied glycosylation (gray) and Man5 (green). *Right,* Structural model created using AF3.

To experimentally characterize these de novo glycoproteins, we expressed each in Expi293F cells, purified them by immobilized metal affinity chromatography, rapidly screened them for the presence of a glycan through PNGase F digestion and gel electrophoresis, and determined glycan composition and occupancy using liquid chromatography-tandem mass spectrometry (LC-MS/MS). From 126 successfully detected de novo glycoproteins, 97 PNGS had >5% occupancy, and on average 94% were complex-type glycans consisting primarily of bi-and tri-antennary structures with high fucosylation and low sialylation typical of HEK-derived glycoproteins (**Figure 1d,e**; **Extended Data Fig. 1**; **Supplementary Data S2**). One design with 31% occupancy carried exclusively the oligomannose-type glycan Man5 in a position predicted to be slightly buried within the protein fold (4_D5; **Figure 1f**). These results establish a scalable framework for designing and characterizing de novo N-linked glycoproteins, while showing that explicit modeling of a Man9 glycan was generally insufficient for encoding oligomannose-type glycans.

### Systematically altering steric restriction controls glycan processing

To systematically test the effects of steric restriction on N-linked glycan processing, we exploited the modular architecture of a de novo twistless helical repeat (THR) protein previously designed to form a 120° corner with a prominent cleft^38^. The THR can be lengthened or shortened by simply adding or removing helix repeat units, enabling precise tuning of the local steric environment of the cleft. We generated a series of four THR glycoproteins (THRgly1–4) in which additional pairs of α-helices progressively enclosed a PNGS located in the cleft, using LigandMPNN to design the glycoprotein sequences (**Figure 2a**). Crystal structures of THRgly1, THRgly2, and THRgly4 revealed that each protein adopted the target fold and steric environment, with Cα RMSDs of 0.87, 0.97, and 0.84 Å, respectively (**Extended Data Fig. 2, Extended Data Table 1**). The glycans were poorly resolved, however, potentially due to flexibility at the exposed site in THRgly1 and low occupancy at the sites in THRgly2 and THRgly4 (**Figure 2b** and **Extended Data Fig. 3**).

**Fig. 2:**
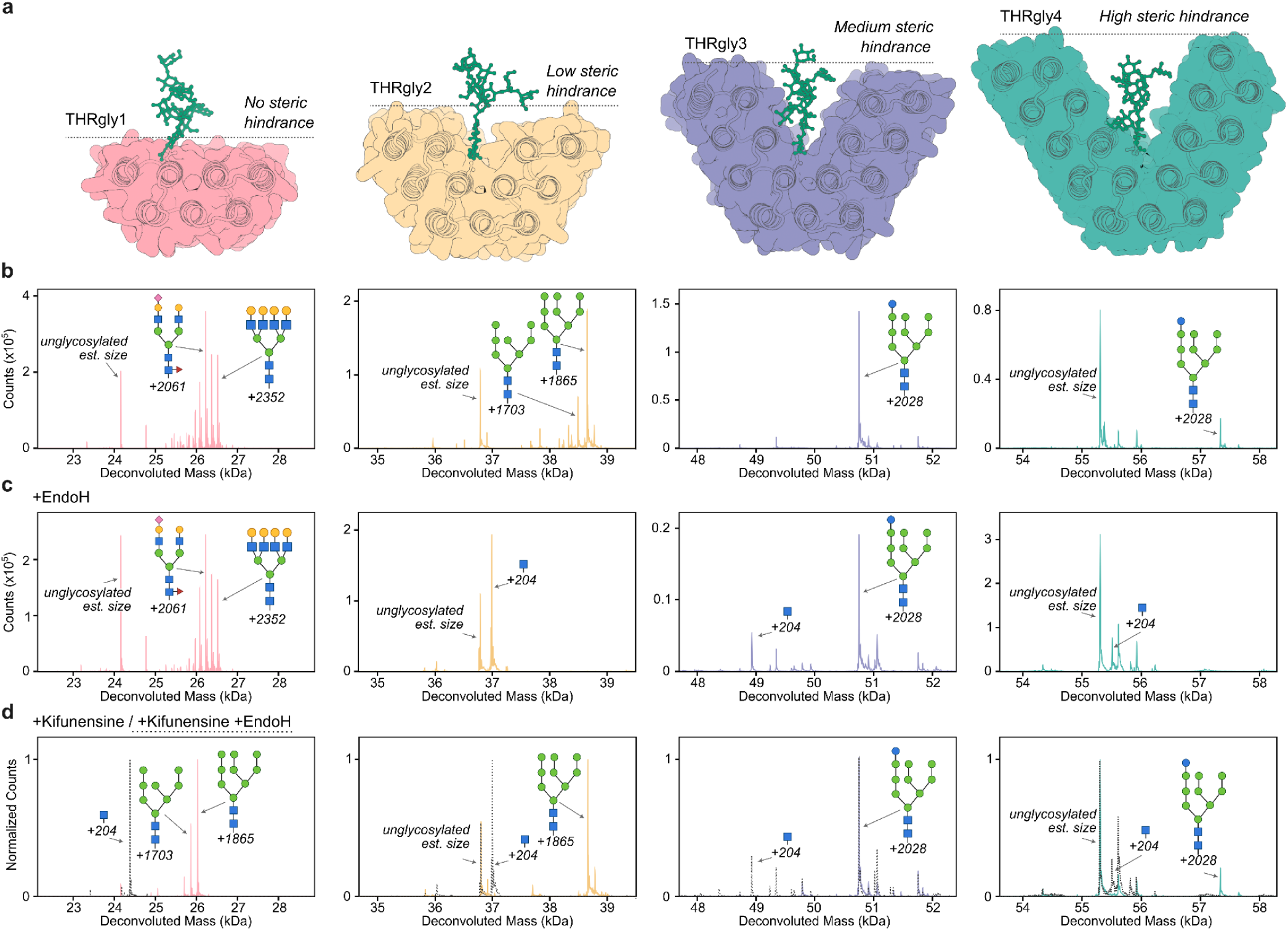
Systematically altering steric restriction controls glycan processing. **a**, AF3-predicted protein with GlycoShape-modeled Man9 glycan (green) for glycoproteins THRgly1-4. **b-d,** Deconvoluted intact mass spectra for analyzing native glycosylation with 1–3 major peaks corresponding to glycosylation shifts labeled. Glycans matching in theoretical molecular weight to the corresponding shift are schematically annotated as follows: mannose residues as green circles, N-acetyl glucosamine residues as blue squares, galactose as yellow circles, sialic acid as pink diamonds, and glucose as blue circles. Unglycosylated sizes are estimated based on predicted MW and labeled, when present. **b:** Native glycosylation. **c:** Native protein digested with Endo H. **d:** Overlaid protein from kifunensine-treated cells (color, solid line) and digested with Endo H (gray, dotted line). Estimated predominant glycans present are shown in MW (Da) differences from predicted unglycosylated size. For peak table see **Supplementary Data S3;** amino acid sequences provided in **Supplementary Data S1**.

**Fig. 3:**
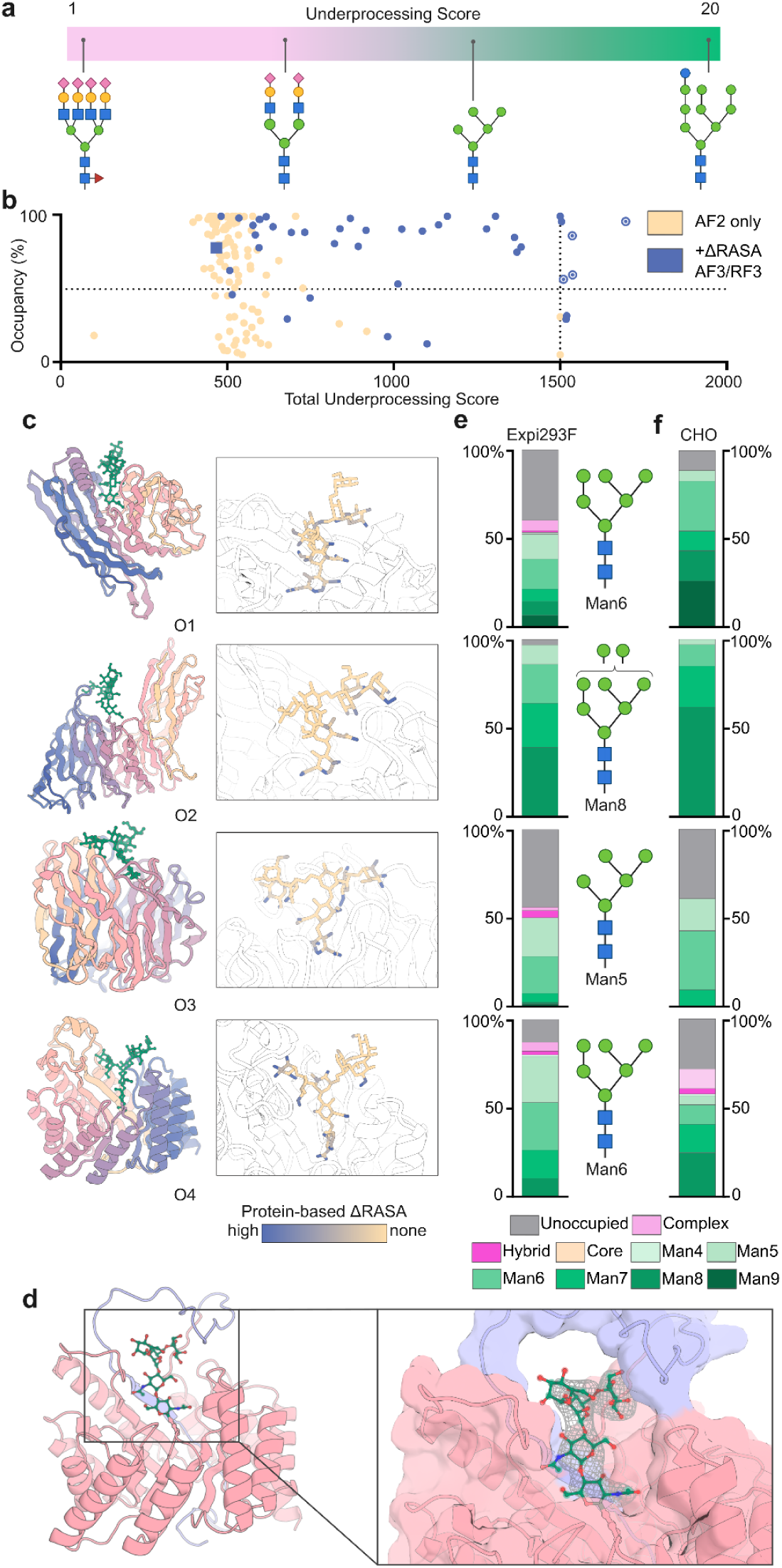
Programming N-glycan processing in de novo glycoproteins. **a**, LC-MS/MS-determined glycans were assigned an ordinal underprocessing score (1-20) based on their position along the N-glycan processing continuum. Lower scores correspond to increasingly processed core/complex-type glycans, while higher scores correspond to oligomannose glycans. Intermediately processed glycans were scored proportionally to their degree of processing. Highest score (20) corresponds to Glc1Man9, representing the least-processed structure. **b**, Resulting score for 140 glycoproteins with >5% occupancy determined from LC-MS/MS data, grouped by first generation designs (light yellow dots) and filtered with ΔRASA (dark blue dots). Oligomannose proteins selected for further analysis highlighted with circled dots, and complex protein control denoted by a square. **c**, *Left,* AF3 predicted structure modeled for de novo oligomannose glycoproteins (O1-4)^30^. Man6 (green) was used during structure prediction. *Right,* Relative accessible surface area (ΔRASA) scoring for individual glycan atoms overlaid on a modeled Man6. High protein-dependent low solvent accessibility shown in dark blue (maximum score=0.51), with glycans lacking any protein-dependent sterics shown in light yellow (minimum score=0). **d**, Resolved crystal structure for O4 at C-alpha RMSD of 2.4Å over residues 51-356. Zoom-in shows partial resolving of modeled Man3. Flexible N-terminal residues (1-50) are colored light blue. **e**, Site specific glycan analysis of Expi293F-secreted glycoproteins determined by LC-MS/MS. Bars represent the abundance of each category of glycan and are color-grouped by compositional isomers: oligomannose-type (green), hybrid (magenta), and complex-type (light pink). Core glycans (light orange) refer to truncated structures smaller than Man3. Unoccupied sites colored in light gray. Full data is available in Supplementary Data S4. **f**, Site-specific glycan analysis of CHO-secreted glycoproteins determined by LC-MS/MS. Bars represent the abundance of each category of glycan and are color-grouped by compositional isomers as in panel **e**.

We used intact mass spectrometry to assess glycan composition in THRgly1–4, as their extreme resistance to denaturation complicated typical peptide-based glycan analysis (**Extended Data Fig. 4**). A range of mass shifts corresponding to various complex glycans was observed for THRgly1, in which the PNGS was exposed on the flat surface of the molecule (**Figure 2b**). By contrast, primarily oligomannose-type glycans were detected on THRgly2–4, in which additional α-helices surrounded some, most, or all of the glycan, respectively. Notably, the predominant glycan in THRgly2–4 increased in size as steric restriction increased: while primarily Man8 and Man9 were observed in THRgly2, THRgly3 and THRgly4 carried glycans with masses consistent with Glc1Man9, an early processing intermediate rarely retained on mature glycoproteins except at exceedingly buried PNGS, such as Asn917 of complement C3 (ref. 39).

To test these assignments, we treated the proteins with Endo H, which specifically cleaves oligomannose-type glycans to leave a single GlcNAc residue detectable as a +204 Da mass shift. In agreement with our mass spectrometry data, this shift appeared in Endo H-treated THRgly2–4 but not THRgly1, the mass spectrum of which appeared relatively unchanged (**Figure 2c**). Digestion was incomplete in THRgly3 and THRgly4, consistent with restricted access to buried glycans, as previously observed for HIV-1 Env^40^. Finally, we produced each of the four proteins in the presence of kifunensine to generate uniformly oligomannose-type glycans regardless of the surrounding protein structure. Intact mass spectra confirmed that each protein, including THRgly1, carried glycans with masses consistent with Man8, Man9, or Glc1Man9, depending on the protein (**Figure 2d**). Endo H treatment produced the +204 Da GlcNAc product in all four cases, further confirming the presence of oligomannose-type glycans in both kifunensine-treated and-untreated THRgly2–4. Together, these data demonstrate that the degree of glycan processing can be controlled by steric restriction encoded in the amino acid sequences of de novo glycoproteins.

### Programming N-glycan processing in de novo glycoproteins

Having established that steric restriction controlled glycan processing in the THRgly repeat-protein architecture, we next sought to develop a general method for encoding oligomannose-type glycans across diverse protein folds. Two methodological advances enabled our approach. First, we incorporated the newly available, glycan-aware structure prediction models RoseTTAFold3 (RF3) and AF3 into our RFdiffusionAA→LigandMPNN→structure prediction pipeline^30,41^. RF3 predictions included a Man9 glycan, whereas AF3 predictions included Man6, the largest glycan accommodated by the server. Second, explicit glycan modeling allowed us to supplement standard structure prediction filters with direct measures of protein-dependent steric restriction. For each design, we calculated the per-atom difference in relative accessible surface area between the glycan in isolation and in the context of the protein (ΔRASA)^42^. After manually reviewing the approximately 5% of designs with the greatest predicted shielding, ranked by either mean per-atom ΔRASA or the number of glycan atoms with nonzero ΔRASA, we selected 82 sequences spanning 24 distinct backbones for expression in Expi293F cells (LC-MS/MS detected amino acid sequences provided in **Supplementary Data S1**).

LC-MS/MS identified a significantly greater fraction of oligomannose-displaying proteins in the 51 detected ΔRASA-filtered set than among the first-generation designs shown in **Figure 1** (20 of 43 vs. 1 of 97 designs exceeding 25% oligomannose content among proteins with >5% glycan occupancy; Fisher’s exact test, p<10^-10^; **Supplementary Data S4**). To compare processing states across the two design sets, we assigned each glycan species a weight inversely proportional to its extent of processing (e.g., Man9 > Man5 > complex-type) and calculated an underprocessing score from the weighted abundances (**Figure 3a and Supplementary Data S5**). Plotting this score against site occupancy allowed us to select one control protein carrying complex-type glycans (C1) and the top four ΔRASA-filtered designs with underprocessing scores >1500 and occupancy >50% (O1–O4) for further characterization (**Figure 3b** and **Extended Data Fig. 5**).

In agreement with the LC-MS/MS data, all four designs were sensitive to Endo H digestion (**Extended Data Fig. 6**). Predicted structures placed the glycans of O1, O3, and O4 within deep pockets or crevices in the protein fold (**Figure 3c**). O2 featured a different structural solution in which a more distant protein ridge shielded a glycan attached at a comparatively shallow site, resulting in a lower ΔRASA score that was similar to control protein C1 (**Extended Data Fig. 7**). Although AF3 predicted all four designs with high confidence, post hoc AF2 analysis indicated that only one design, O2, would have passed our original filters (**Extended Data Table 2**). This result made intuitive sense, as AF2 modeled each sequence without the glycan and would likely interpret the deep glycan-housing pockets in O1, O3, and O4 as large, unoccupied cavities. Circular dichroism spectra were consistent with the designed secondary structure of each protein and showed that only O3 exhibited a clear thermal unfolding transition, at 79 °C, while the other proteins did not show a cooperative transition below 95 °C (**Extended Data Fig. 8**). A crystal structure of O4 at 3.9 Å resolution closely matched the design model over most of its length (Cα RMSD of 2.4 Å over residues 51–356), although the N-terminal 50 residues deviated from the design to form a lid-like structure over the glycan (**Figure 3d**). Unlike the THRgly structures, the O4 map contained electron density consistent with the first five monosaccharide residues of the glycan within this pocket. This observation is consistent with the surrounding protein restricting glycan conformational mobility as well as processing.

The precise distribution of oligomannose-type glycans in each glycoprotein varied, with O2 enriched for Man7–9, while O1, O3, and O4 carried more processed species, including Man5 and Man6 (**Figure 3e**). To determine whether these processing states persisted in another expression system, we produced the designs in Chinese hamster ovary (CHO) cells. All four oligomannose-type glycoproteins had glycan compositions comparable to those observed in Expi293F cells by LC-MS/MS (**Figure 3f** and **Supplementary Data S6**). The control glycoprotein C1 likewise carried a similar distribution of complex-type glycans in both cell lines (**Extended Data Fig. 5** and **Supplementary Data S6**). These results are consistent with local protein structure, rather than the expression host, being the primary determinant of glycan processing state. Together, these data establish a general method for programming both the location and composition of N-linked glycans in de novo proteins.

### De novo oligomannose-type glycoproteins inhibit UPEC adhesion and engage innate immune lectins

Having established that designed protein structure can program oligomannose-type glycan composition, we next asked whether the resulting glycans remained accessible and could mediate biological activity. Uromodulin (Tamm-Horsfall protein), the most abundant protein in urine, provided a useful precedent as an unusual mature human glycoprotein bearing a functionally exposed oligomannose-type glycan^43^. Located at Asn275, this glycan acts as a soluble receptor decoy for FimH, the mannose-binding adhesin used by uropathogenic *E. coli* (UPEC) to attach to uroepithelial cells during colonization^28,29^. Because FimH preferentially recognizes smaller oligomannose species, particularly Man5 (ref. ^27^), we tested whether the Man5-rich O3 and O4 glycoproteins could compete for FimH binding.

We preincubated high-affinity FimH pili with each glycoprotein and measured residual binding to RNase B, a Man5-bearing glycoprotein that binds FimH strongly^44^. O3 reduced FimH binding by 75 ± 5% (mean ± SD, Dunnett’s test p=0.0205), while O4 reduced binding by 44 ± 14% (p=0.0041), followed by O2 at 41 ± 20% (p=0.0253) (**Figure 4a** and **Supplementary Data S7**). The complex-type control protein C1 did not significantly reduce binding (p=0.726). The differing activity of O3 and O4 despite carrying similar proportions of Man5 suggests that local glycan presentation—including potential shielding by the lid-like loop observed in the O4 crystal structure (**Figure 3d**)—may also influence FimH recognition. The comparatively weak activity of O2, which was enriched for the larger oligomannose species Man7–Man9, was consistent with the established preference of FimH for smaller oligomannose structures.

**Fig. 4:**
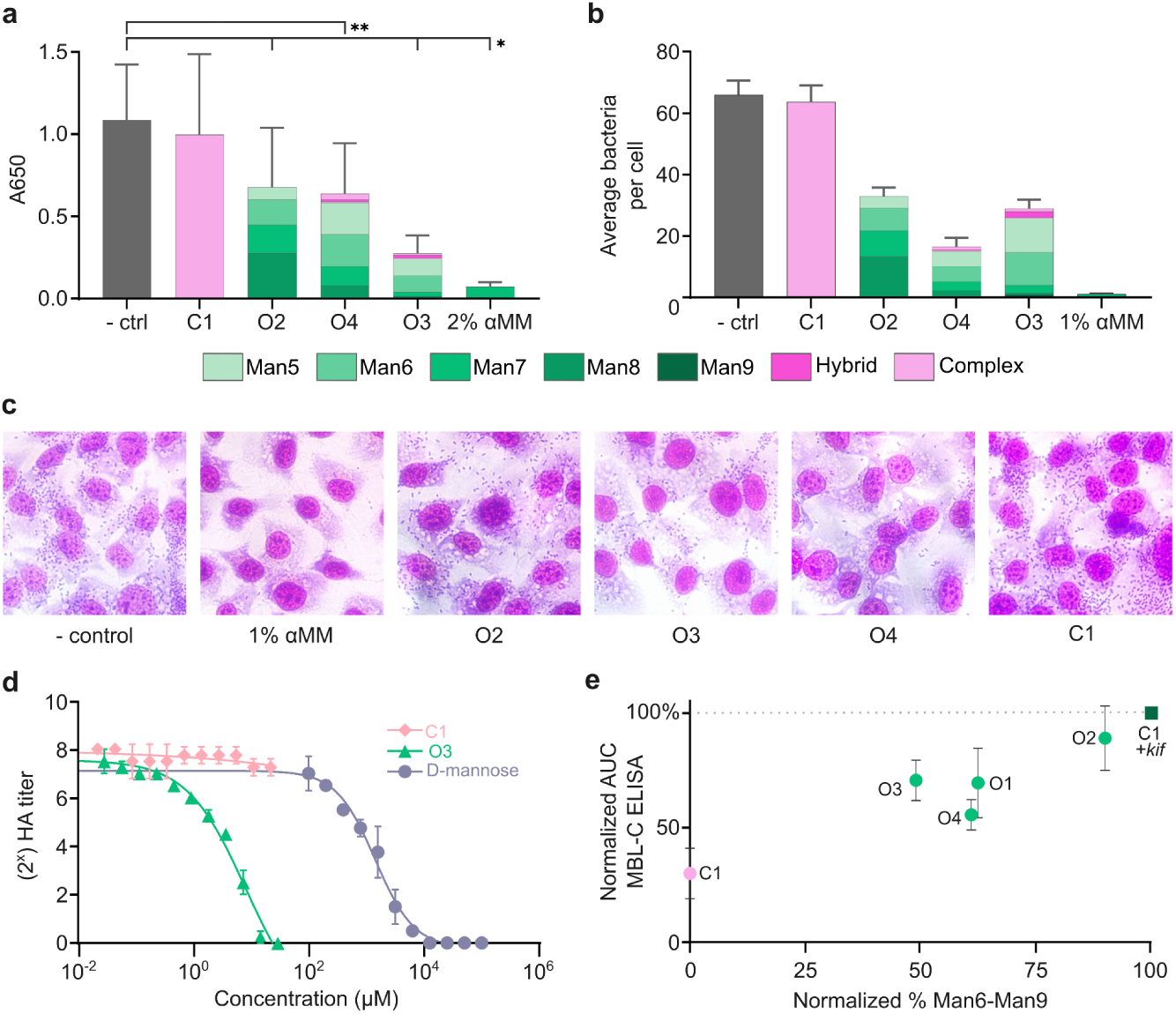
De novo oligomannose-type glycoproteins inhibit UPEC adhesion and engage innate immune lectins. **a**, RNase B competition assay. Proteins (20 μg), 2% α-methyl-D-mannopyranoside, or 0.1% BSA in PBS were pre-incubated with high-affinity FimH and added to immobilized RNase B. Bound FimH was detected by anti-fimbrial antibody (pAb280). Data are mean ± SD of raw absorbance of four independent experiments. Repeated-measures one-way ANOVA with Geisser-Greenhouse correction (p=0.013, n=4) and Dunnett’s multiple comparisons test versus 0.1% BSA in PBS (*p<0.05, **p<0.01). De novo protein bars colored in chart by LC-MS/MS-based determination of glycan composition for occupied sites (Figure 3e). **b**, Average bacterial adherence per T24 cell (n=20 from a single experiment; error bars show SEM across cells) after pre-incubation of bacterial cells with 20 μg/mL de novo proteins, 2% FBS, or 1% α-methyl-D-mannopyranoside. **c**, Representative images of the data summarized in **b**: T24 bladder cells after infection with pre-incubated UPEC and stained with eosin Y followed by Hema3. **d**, HA titration using UPEC strain UTI89 and guinea pig erythrocytes, with serial dilutions of de novo protein or D-mannose (n=2). **e,** MBL-C ELISA, quantifying area under the curve (AUC). De novo proteins were serially diluted and allowed to bind to immobilized MBL-C. Complex glycoprotein C1 as well as C1 treated with kifunensine served as controls. Data are normalized to kifunensine control, and shown as mean ± SD of two independent experiments. Normalized % Man6-Man9 are calculated as relative to total occupancy.

We next asked whether the de novo glycoproteins could inhibit UPEC attachment to T24 human epithelial bladder cells, a physiologically relevant model of urinary tract infection. O2, O3, and O4 reduced UPEC attachment by 50%, 56%, and 75% relative to the negative control, whereas C1 had no detectable effect at 3% reduction of control adherence (**Figure 4b,c** and **Supplementary Data S8**). O3 also inhibited red blood cell agglutination by *E. coli* UTI89 with an EC_50_ of 8 μM [CI: 4.3–84.5 μM, based on log_2_ hemagglutination (HA) titer], 100-to 1000-fold lower than the concentration of mannose required for inhibition^44^ (**Figure 4d** and **Supplementary Data S9**).

Finally, we tested whether the designed glycans could engage mammalian MBL (mouse MBL-C), a soluble component of the complement system that preferentially recognizes mannose-rich glycans^45–48^. Consistent with this preference, we found MBL binding to kifunensine-treated C1 was approximately threefold higher than to untreated C1 bearing complex-type glycans (**Figure 4e** and **Supplementary Data S10**). All four de novo glycoproteins bound MBL more strongly than untreated C1. In contrast to FimH, MBL bound most strongly to O2, which predominantly displayed Man8 and reached 79–99% of the binding observed for the kifunensine-treated control in two independent experiments. Together, these results demonstrate that de novo amino acid sequences can be designed to encode glycan-driven, glycoform-dependent biological activity.

## DISCUSSION

Here we establish local protein structure as a programmable determinant of N-linked glycan processing in de novo glycoproteins. In a systematic series of twistless helical repeat proteins, progressive enclosure of an N-linked glycan shifted its composition toward increasingly underprocessed oligomannose-type species. We then generalized our design approach across diverse de novo folds and found that the glycan profiles of selected designs were similar in two mammalian expression systems. Unlike approaches that globally alter cellular glycosylation^9,10^, our method encodes a preferred processing state in the structure—and therefore the sequence—of the designed protein itself. This property could be useful in recombinant glycoprotein manufacturing and, if maintained upon expression in vivo, in genetic approaches to biologics delivery such as mRNA^49^. More broadly, our approach provides a route to making both glycan position and composition programmable features of protein-based medicines and materials.

Controlling glycan processing state required design and prediction methods that explicitly model glycans along with proteins. Our initial pipeline used RFdiffusionAA and LigandMPNN to design around a modeled glycan, but relied on AF2 structure prediction to filter the resulting sequences. Among 97 first-generation designs with >5% glycan occupancy, only one exceeded 25% oligomannose content. Combining glycan-aware RF3 and AF3 predictions with an explicit measure of glycan burial during filtering substantially increased the fraction of designs that were experimentally confirmed as oligomannose-type glycoproteins. However, while generally biasing glycan processing toward underprocessed glycoforms, this approach did not specify a single, homogeneous glycan structure. The Glc1Man9 glycans observed in THRgly3 and THRgly4 indicate that more extensive restriction is possible, and the extreme denaturation resistance of these proteins motivates further study of pocket geometry and conformational rigidity as determinants of processing outcomes. Such studies, combined with continued improvements in all-atom design and prediction methods, should enable increasingly fine control over site-specific glycan composition^42,50^.

Despite their programmed steric restriction, the glycans in our de novo glycoproteins remained accessible for biological recognition, enabling inhibition of FimH-dependent UPEC adhesion and glycan composition-dependent binding of MBL. Engagement of both FimH and MBL, which have contrasting glycoform preferences, showed how programmed glycan composition can direct biological activity without a designed protein-protein interface. Looking forward, the MBL-binding activity of O2 motivates combining programmable glycan composition with de novo nanoparticle vaccine design, as previous studies established roles for MBL and complement in the trafficking and efficacy of glycosylated nanoparticle immunogens^47,51–53^. For FimH, the different blocking activities of O3 and O4 despite their similar Man5 content suggest that the surrounding scaffold also shapes target engagement. Future designs could therefore co-optimize glycan-and protein-mediated contacts; the anti-FimH antibody mAb475 provides a natural precedent, using a high-mannose glycan to occupy the FimH mannose-binding pocket while adjacent residues form additional contacts^54^. More broadly, this strategy could be extended beyond UPEC to generate glycoprotein receptor decoys for other glycan-binding pathogens, such as influenza viruses (sialic acid) or noroviruses (histo-blood group antigens)^55^. Glycans play numerous roles in biology, ranging from stabilizing individual proteins to forming structural polymers in crustacean shells and wood^2,56^. Extending sequence-encoded control to other glycan classes could therefore broaden the functions accessible to designed protein medicines and materials.

### Data, Materials, and Software Availability

Data deposition, atomic coordinates, and structure factors for all crystal structures have been deposited in the Protein Data Bank, http://www.rcsb.org (PDB IDs: 39BF, 39BG, 39BH, and 39BI). Raw files associated with LC-MS/MS analysis of glycoproteins generated in this study are deposited at massive.ucsd.edu under dataset MSV000103421. AF3-Server design models for all LC-MS/MS-detected and THR glycoproteins with a modeled Man6 can be downloaded using the following stable link: https://files.ipd.uw.edu/pub/mlunn/AF3_predictions.zip

## Author Contributions

M.C.LH., J.D.A., M.C., D.B., and N.P.K. conceived the study. R.K. and W.A. developed RFdiffusionAA and RASA-based filtering. M.C.LH., J.D.A. designed proteins, expressed and characterized glycoproteins. M.C.LH. performed MBL ELISAs. J.D.A. performed LC-MS/MS. A.M. carried out RNase B inhibition and T24 assays. X.L. and C.M. performed mass spectrometry. J.P. performed HA titration and E.L. analyzed data. A.K., H.N., J.M., E.J. and A.B. conducted crystallographic experiments, data processing, structure solution, and final refinement. E.S. supervised RNaseB and T24 assays. S.J.H. supervised HA titration. M.C. supervised LC-MS/MS analysis. M.C.LH., J.D.A., and N.P.K. wrote and revised the manuscript, with input from all co-authors.

## Acknowledgements

The authors thank Kandise VanWormer, Hernan Nunez-Ortega, and Eryn Weston for laboratory and administrative support; Suna Cheng, Craig Dobbins, and Adri Tran-Peterson for tissue culture assistance; Luki Goldschmidt and Patrick Vecchiato for maintaining computing infrastructure at the Institute for Protein Design; and Cyrus Haas for general support. This work was supported by the Gates Foundation (INV-069938 to D.B. and N.P.K.), the National Institute of Allergy and Infectious Diseases (U19AI181881 to D.B. and N.P.K., U19AI157797 to S.J.H.), the Advanced Research Projects Agency for Health (ARPA-H; APECx Program Award No. 1AY1AX000036-01 to D.B. and N.P.K.), and the National Science Foundation Graduate Research Fellowships Program Award (DGE-2140004 to M.C.LH.). This work is based upon research conducted at the Northeastern Collaborative Access Team beamlines, which are funded by the National Institute of General Medical Sciences from the National Institutes of Health (P30 GM124165). The Eiger 16M detector on the 24-ID-E beam line is funded by a NIH-ORIP HEI grant (S10OD021527). This research used beamtime awards (DOI: https://doi.org/10.46936/APS-190193/60014259) from the Advanced Photon Source, a U.S. Department of Energy (DOE) Office of Science User Facility operated for the DOE Office of Science by Argonne National Laboratory under Contract No. DE-AC02-06CH11357. This research used resources 17-ID-1/17-ID-2 of the National Synchrotron Light Source II, a U.S. Department of Energy (DOE) Office of Science User Facility operated for the DOE Office of Science by Brookhaven National Laboratory under Contract No. DE-SC0012704. This publication resulted from data collected using beamtime obtained through NECAT BAG proposal #317877. The Center for BioMolecular Structure (CBMS) is primarily supported by the National Institutes of Health, National Institute of General Medical Sciences (NIGMS) through a Center Core P30 Grant (P30GM133893), and by the DOE Office of Biological and Environmental Research (KP1605010).

## METHODS

### Computational redesign of THRgly de novo proteins to display oligomannose glycans

To design protein backbones, previously described protein building blocks that take stable, higher order structures were redesigned and truncated^38^. ProteinMPNN was run with 500 sequences per target and per-position probability distributions were saved for analysis^57^. Positions predicted to favorably accommodate an N with a downstream sequon-compatible T or S were selected as candidates for mutagenesis. Man9 glycan was modeled on the mutated asparagine residue using GlycoShape, and glycan-aware LigandMPNN was used to redesign surrounding side chains^31,34^. To improve secretion and solubility, SolubleMPNN was used as a recycling step to mutate exposed hydrophobic residues^57^. AF2 was used to predict structures without glycans and high confidence structures were selected for further characterization (pLDDT>90, pAE <5, Cα-RMSD <2Å)^35^. Following the release of AF3, we then re-predicted these structures with a Man6 for comparison to crystal structure models^30^.

### De novo glycoprotein design

De novo glycoproteins were generated using RoseTTAFold diffusion All-Atom to generate backbones^32^. As inputs to RFdiffusionAA, a diverse panel of PNGS or asparagines alone was selected across different secondary structures and origins (viral, human, and designed bacterial). Structures were isolated and GlycoShape was used to attach a single Man9 glycan to the asparagine residue [PDB: 6vxx^58^ (8 sites; N61, N164, N235, N331, N603, N709, N717, N1134), PDB: 7pfp^29^ (3 sites; N232, N396, N513), I53-50A-3gly^59^ (3 sites; N45, N75, N96)]. An isolated peptide fragment from the crystal structure of OST-A’s binding pocket was also used, with the aim of increasing overall occupancy rates, and 5 glycan conformations were modeled to increase diversity using GlycoShape [PDB:6S7O (1 site; N3)^12^]. RFdiffusionAA generated a total of ∼150,000 backbones ranging from 300–500 amino acids using 50–75 diffusion steps, guided by a ligand contact potential (ligand_ncontacts, weight=0.5–1, guide scale=2, quadratic decay) to bias folds towards glycan-protein contacts.

For all sets of designs, LigandMPNN was used to design optimal soluble sequences with a glycan included as ligand context and SolubleMPNN or EnhancedMPNN as a recycling step to increase sequence diversity in the ΔRASA-based campaign (below)^31,57,60^. A default sampling temperature of 0.1 and 5–8 sequences per backbone was used across models, with each NXT or NXS motif fixed and cysteines omitted. For a small selection of proteins, we mutated the motif isolated from the peptide fragment from the crystal structure of OST-A’s binding pocket to introduce a point mutation at the-2 position shown to increase occupancy (FANATAA)^12,37,61^.

An initial set of designs was generated without any ΔRASA-based metrics or conditioning. As AF2 was the only structure prediction model available at the time, designed sequences were predicted using AF2 in single-sequence mode and selected for experimental characterization based on high confidence (pLDDT>80, pAE <10, Cα-RMSD < 3 Å).

Following the release of all-atom prediction models, a second set of designs was generated using the same inputs as the RFdiffusionAA pipeline, but incorporating RASA-based conditioning during generation (via a pre-release version) intended to bias sampling towards specific burial values. Post hoc analysis revealed that this conditioning had no measurable effect, and we therefore treat these designs as equivalent to unconditioned generations in analyses. These designed sequences were predicted using RF3 with default weights and a modeled Man9, or AF3 Server (alphafoldserver.com) with an attached Man6 (due to server size limits)^30,41^. Designs were selected for experimental characterization based on a confidence filter for RF3 or AF3 (pLDDT>75, pAE <15, Cα-RMSD < 8 Å to design model).

For this ΔRASA-based set, glycan burial was assessed post-prediction using a relative accessible surface area (RASA), using a 1.4 Å probe (Shrake-Rupley, biopython) normalized per atom to the theoretical maximum accessibility of an isolated atom, 4π(r + r_probe)^2^, as previously described^42,62^. RASA was computed twice for each structure: once for the glycan atoms in the context of the protein, and once for the glycan in isolation to avoid burial attributable to intramolecular glycan-glycan contacts. ΔRASA was calculated as the absolute difference between these values, where higher values represent greater occlusion by protein structure. Designs were ranked by average ΔRASA score together and the number of atoms with a ΔRASA of nonzero (contact). High-ranking predictions by either summary (approximately the top 5%) were manually inspected to select final designs for experimental characterization.

### Experimental Screening and Protein Production

Glycoprotein designs were synthesized as genes (eBlocks, Integrated DNA Technologies) with compatible BsaI overhangs for cloning into a modified pCMV/R expression vector with a Carbenicillin resistance gene or cloned into pCMV/R expression vector between restriction sites XbaI and AvrII with a Kanamycin resistance gene by GenScript. The final sequence contained an N-terminal secretion signal sequence with a C-terminal 6xHis affinity tag.

Signal sequences were either derived from the modified bovine prolactin or mu-phosphatase signal peptide (MDSKGSSQKGSRLLLLLVVSNLLLPQGVLA or MGILPSPGMPALLSLVSLLSVLLMGCVAETGT). C-terminal tags included either a Myc-tag (EQKLISEEDLHHHHHH), TEV-cleavage site (ENLYFQGHHHHHH), or WELQut cleavage site (WELQEGSHHHHHH).

Golden gate reaction mixtures or plasmids were transformed into NEB5-alpha (New England Biolabs C2987). Transformations were then grown in 2-200mL cultures in autoclaved LB media with 50 µg/mL Carbenicillin or Kanamycin at 37°C at 250 rpm. Plasmid DNA was then harvested using Qiagen Spin miniprep kits (Qiagen 27104), ZymoPURE miniprep kits (Zymo D4210), AmMag™ Quatro Maxi (GenScript), or Plasmid Plus maxi kit (Qiagen 12963), and 1µg purified DNA was transfected per 1mL of 3.0×10^6^ Expi293F cells (ThermoFisher A14527) using 3µg of PEI-MAX (Polysciences). For initial screening, cells were shaken at 37°C and 8% CO2 with shaking (orbital shaker, 250 rpm) in TPP TubeSpin bioreactor tubes (Millipore Sigma Z761028) at 10mL, and for large scale protein production volumes ranged from 0.5-3 L at slower speeds (orbital shaker, 150 rpm). After 3 days, supernatant was harvested by centrifugation for 5 minutes at 4500 rpm followed by the addition of PDADMAC (Sigma Aldrich), a second centrifugation for 5 minutes at 4500 rpm, and sterile filtration. For kifunensine-treated cells, 46.5 μL of 1 mg/mL was added per 10mL cells, for a final concentration of 20 µM.

To test alternative cell lines, recombinant proteins were produced using TurboCHO™ (GenScript) expression system at a 4mL scale and supernatants were tested alongside Expi293F recombinant proteins.

Supernatant was adjusted to 50 mM Tris (pH 8.0) and 350 mM NaCl. Nickel Sepharose Excel (Millipore Sigma GE17-3712-02) was added to supernatant and placed on a rocking platform at 4°C overnight. Resin was collected in a gravity column and washed with Wash Buffer (50 mM Tris, 500 mM NaCl, 30 mM Imidazole). Protein was eluted with Elution Buffer (50 mM Tris, 500 mM NaCl, 300 mM Imidazole). For further purification, eluates were applied to a Superdex 200 HPLC or SEC column (Cytiva) equilibrated with AKTA buffer (25 mM Tris, 150 mM NaCl). Peaks corresponding to designs were selected based on volume.

To digest glycoproteins, Endoglycosidase H and Peptide-N-Glycosidase F were used according to manufacturer’s instructions (NEB P0702L, P0704L). Proteins were run on 4–20% Stain Free Criterion TGX gels (BioRad 5678095) for 30-60 minutes at 180V. Gels were stained using GelCode Blue Stain Reagent (ThermoFisher 24592).

### Circular Dichroism

Proteins were diluted in 25 mM Tris, 150 mM NaCl buffer to 0.5–1 mg/mL. Samples were run on a JASCO-15000 equipped with a temperature controlled multi-cell holder, and data were analyzed using GraphPad Prism software. Full wavelength scans were performed at 25, 35, 45, 55, 65, 75, 85, and 95 °C at a heating rate of 2 °C/min. Readings at 217 or 222 nm were recorded every 1 °C.

### Intact Mass Spectrometry

Samples were prepared at 0.5–1 mg/mL in 25 mM Tris, 150 mM NaCl and free cysteines were reduced using 20 mM TCEP. Samples were desalted using Zeba 7K MWCO. To identify the molecular mass of each protein, intact mass spectra were obtained via reverse-phase LC/MS on an Agilent G6230B TOF on an AdvanceBio RP-Desalting column from 10% acetonitrile in 0.1% formic acid to 95% acetonitrile in 0.1% formic acid over the course of 2 minutes, and subsequently deconvoluted by way of Bioconfirm using a total entropy algorithm from m/z 600 to 2500.

### Liquid chromatography-mass spectrometry analysis of glycoproteins

Samples were denatured in 6 M urea, 50 mM Tris/HCl, pH 8.0, for 2 h, buffer-exchanged into 50 mM Tris/HCl, pH 8.0 using Vivaspin columns (10 kDa), and digested overnight with trypsin-LysC (Promega), chymotrypsin or alpha-lytic protease (Sigma) at an enzyme:protein ratio of 1:30 (w/w). (Glyco)peptides were desalted on Oasis HLB µElution plates (Waters), resuspended in 0.1% formic acid, and analysed by LC-ESI-MS on a Vanquish Neo UHPLC system coupled to an Orbitrap Eclipse Tribrid (Thermo Fisher Scientific) with stepped HCD fragmentation. Samples were first run on a 30 min high-throughput gradient; those with insufficient glycopeptide coverage were re-analysed on an extended 180 min gradient.

For the 30 min method, peptides were trapped on a 300 µm × 0.5 cm cartridge and separated on a 75 µm × 5.5 cm μPAC High throughput column (50 °C) at 1.0 µL/min with A = 0.1% formic acid and B = 80% acetonitrile / 0.1% formic acid, using a gradient that rose from 5% B at 0 min to 50% B at 25 min and 95% B at 26 min, held at 95% B until 27 min, then returned to 5% B from 28 to 32 min for re-equilibration. MS1 spectra were acquired in the Orbitrap at 240,000 resolution over 200–2000 m/z, and MS2 spectra were acquired in the Orbitrap at 30,000 resolution using stepped HCD at 15, 30 and 45% NCE. Samples re-run on the 180 min method were separated on a μPAC Neo column (180 µm × 110 cm) using a 180 min linear gradient of 4–32% acetonitrile in 0.1% formic acid, followed by a 15 min wash at 80% acetonitrile in 0.1% formic acid. MS acquisition parameters were as described for the 30 min method.

Raw files from both methods were processed in Byos v5.5 (Protein Metrics) and searched against the Protein Metrics 305 N-glycan library with precursor and fragment mass tolerances of 4 and 10 ppm, respectively, and a 1% false discovery rate (FDR). Glycopeptides were validated manually, requiring the expected b/y fragment ions together with oxonium ions for the assigned glycan. The relative abundance of each glycan at each site, and the proportion of unoccupied sites, were quantified from summed extracted ion chromatogram (XIC) areas across all charge states of a given glycopeptide. Glycans were classified by composition: HexNAc(2)Hex(9−3) as Man9–Man3, fucosylated forms as FM, complex-type glycans by HexNAc count and fucosylation, and truncated species smaller than Man3 as core glycans; isomeric glycans sharing a composition were not distinguished.

### Inhibition of *E. coli* adhesion to RNase B

Immulon 4HBX 96-well plates were coated with 20 µg/mL RNase B and quenched with 0.1% BSA in PBS. Bacterial fimbriae expressing the high mannose-binding FimH–FocH adhesin hybrid were adjusted to a concentration of 0.2 mg/mL, mixed 1:1 with the tested glycoproteins at concentrations of 20 µg/mL, and incubated at room temperature for 30 min. As a control, fimbriae were incubated with 0.1% BSA alone or with the addition of 2% methyl-α-D-mannopyranoside. After the incubation, all samples were added to the wells coated with RNase B and incubated for 45 min at 37 °C, followed by two washes with PBS. Rabbit anti-fimbrial polyclonal antibodies (Pab280) were diluted 1:1000 and added to the wells for 50 min, followed by extensive washing. Antibody binding was detected with goat anti-rabbit IgG antibodies conjugated to horseradish peroxidase, incubated for 30 min. After extensive washing, a colorimetric reaction was developed by adding 3,3′,5,5′-tetramethylbenzidine (TMB) to the wells, and absorbance was read at 650 nm.

### *E. coli* bladder cell adhesion assays

Bacterial cell suspensions (50 μL of 10^9^ CFU/mL) were mixed 1:1 with McCoy’s 5A medium with 2% heat-inactivated fetal bovine serum, without added inhibitors or with 1% α-methyl-D-mannopyranoside or 20 μg of the glycoproteins and were added to monolayers of T24 human bladder epithelial carcinoma cells grown in 12-well plates. Bacteria were allowed to interact with the cells for 2 h at 37 °C in 5% CO_2_. The cells were washed six times with PBS, fixed with cold 100% methanol for 1 min, air dried and stained with eosin Y followed by Hema3, both dyes diluted 1:2 with water.

### Hemagglutination inhibition assay

Erythrocyte hemagglutination inhibition assays using *E. coli* UTI89 were performed as previously described^63^. Briefly, serial dilutions of the tested compounds and proteins were prepared in microtiter plates, with each row corresponding to a single inhibitor concentration. The first well of each row received 25 μL of UTI89 bacterial suspension in 1× PBS at a starting OD₆₀₀ of 10.0, which was then serially diluted across the row. Subsequently, 25 μL of guinea pig erythrocytes (OD₆₄₀ = 2.0; Colorado Serum Company) suspended in 1× PBS was added to each well, resulting in a final assay volume of 50 μL per well. Plates were incubated overnight at 4 °C to allow hemagglutination to develop. The HA titer at each inhibitor concentration was defined as the highest bacterial dilution that produced visible agglutination of the erythrocytes from a single experiment.

### Recombinant Mannose-binding Lectin ELISA

Selected glycoproteins were coated for two replicates directly onto MaxiSorp plates in 50µL PBS at 1 µg/mL. Plates were incubated at 4°C overnight, then washed with PBS. Plates were blocked in PBS containing 1% BSA and 0.1 M CalCl_2_ for 2 hours at room temperature. Plates were washed four times in PBS containing 0.1% Tween 20. Recombinant mouse MBL-C (2208-MB-050/CF, R&D) was prepared at two-fold serial dilutions beginning at 50 µg/mL. 50µL of each concentration was added to the plate and incubated for one hour at 37°C. Plates were washed four times in PBS containing 0.1% Tween 20. 50µL of anti-MBL antibody (14D12, Abcam) at 2 µg/mL in blocking buffer was added to each well and incubated at room temperature for 1 hour. Plates were washed four times in PBS containing 0.1% Tween 20. 50µL of a 1:5000 dilution of anti-rat IgG:HRP (7077S, Cell Signaling Technologies) in blocking buffer was added to each well, then the plate was incubated for 1 hour at room temperature. Plates were washed four times in PBS containing 0.1% Tween 20. 50µL of TMB (1-Step Ultra TMB, Thermo Fisher) was added and plates were developed until the lowest dilution wells were visibly showing signal. Reaction was stopped with the addition of 1N hydrochloric acid. Plates were read at 450 nm with a reference wavelength of 540 nm. Duplicate wells were averaged. AUC was calculated in GraphPad Prism.

### Statistical Analyses

Analyses were performed in GraphPad Prism v11.0.0. For comparison between design sets, the fraction of proteins with >25% oligomannose among those with >5% occupancy was compared between sets by two-sided Fisher’s exact test. For the RNase B competition assay, raw absorbance (650 nm) from four independent experiments was analyzed by repeated-measures one-way ANOVA with Geisser-Greenhouse correction (sphericity not assumed), followed by Dunnett’s multiple comparison test against 0.1% BSA in PBS. Adjusted P values reported. For HA titers (log_2_), two replicate wells per concentration were fit to a variable-slope-dose-response model. EC50 values are reported with 95% confidence intervals. For the MBL ELISA, area under the curve (AUC) was calculated for each independent experiment by the trapezoidal rule from absorbance (450 nm) versus MBL concentration (µg/mL), with a baseline of zero and normalized to the AUC of kifunensine-treated C1 (100%) from the same experiment. Data are mean ± SD of two independent experiments.

### Crystallographic Structure Determination

All crystallization experiments were conducted using the sitting drop vapor diffusion method. Crystallization trials were set up in 200 nL drops using the 96-well plate format at 20 °C. Crystallization plates were set up using a Mosquito LCP from SPT Labtech, then imaged using UVEX microscopes and UVEX PS-256 from JAN Scientific. Diffraction quality crystals formed in 0.2 M Calcium chloride, 0.1 M HEPES, pH 7, and 20% (w/v) PEG 6000 for THRgly4, in 0.1 M Sodium citrate tribasic dihydrate, pH 5.6, 2.0 M Ammonium sulfate, and 0.2 M Potassium sodium tartrate tetrahydrate for THRgly1 and in 0.1 M Sodium cacodylate, pH 6.5, 40% (v/v) MPD, 5% (w/v) PEG 8000 for THRgly2, and in 0.12 M ethylene glycols (Diethylene glycol, triethylene glycol, tetraethylene glycol, pentaethylene glycol); 0.1 M Buffer System 1 (Imidazole:MES), pH 6.5; 30% v/v Precipitant Mix 1 (20% PEG 500 MME, 10% PEG 20,000) for O4.

Diffraction data was collected at the Advanced Photon Source beamline on 24-ID-E and National Synchrotron Light Source II on FMX/AMX. X-ray intensities and data reduction were evaluated and integrated using XDS^64^ for THRgly1 & THRgly4 and xia2/DIALS^65,66^ for THRgly2 & O4 and merged/scaled using Pointless/Aimless in the CCP4 program suite^67^. Structure determination and refinement starting phases were obtained by molecular replacement using Phaser^68^ for THRgly1 & THRgly4, DIMPLE^69^ for THRgly2, and MRBUMP^70^ for O4 using the designed model for the structures. For THRgly1 & THRgly4 following molecular replacement, the models were improved using phenix.autobuild^71^; efforts were made to reduce model bias by setting rebuild-in-place to false, and using simulated annealing. Structures were refined in Phenix^71^. Model building was performed using COOT^72^. The final model was evaluated using MolProbity^73^. For THRgly2 and O4, iterative model building and corrections were performed manually using COOT^72^ following molecular replacement and subsequent structure refinement was performed with CCP4 Refmac5^74^and CCP4 Cloud Refmacat^75,76^. Initial refinement was conducted using BUSTER^77^ to rapidly fix Ramachandran, rotamer, and density fit outliers, refining to convergence. PDB-REDO^78^ was used to assess the model quality in between refinements and to fix any rotamer and density fit outliers. Final refinement was carried out in Phenix^71^, with manual model building performed in COOT^72^.The model quality was evaluated using MolProbity^73^. Data collection and refinement statistics are recorded in **Extended Data Table 1**.

## EXTENDED DATA

**Extended Data Figure 1:**
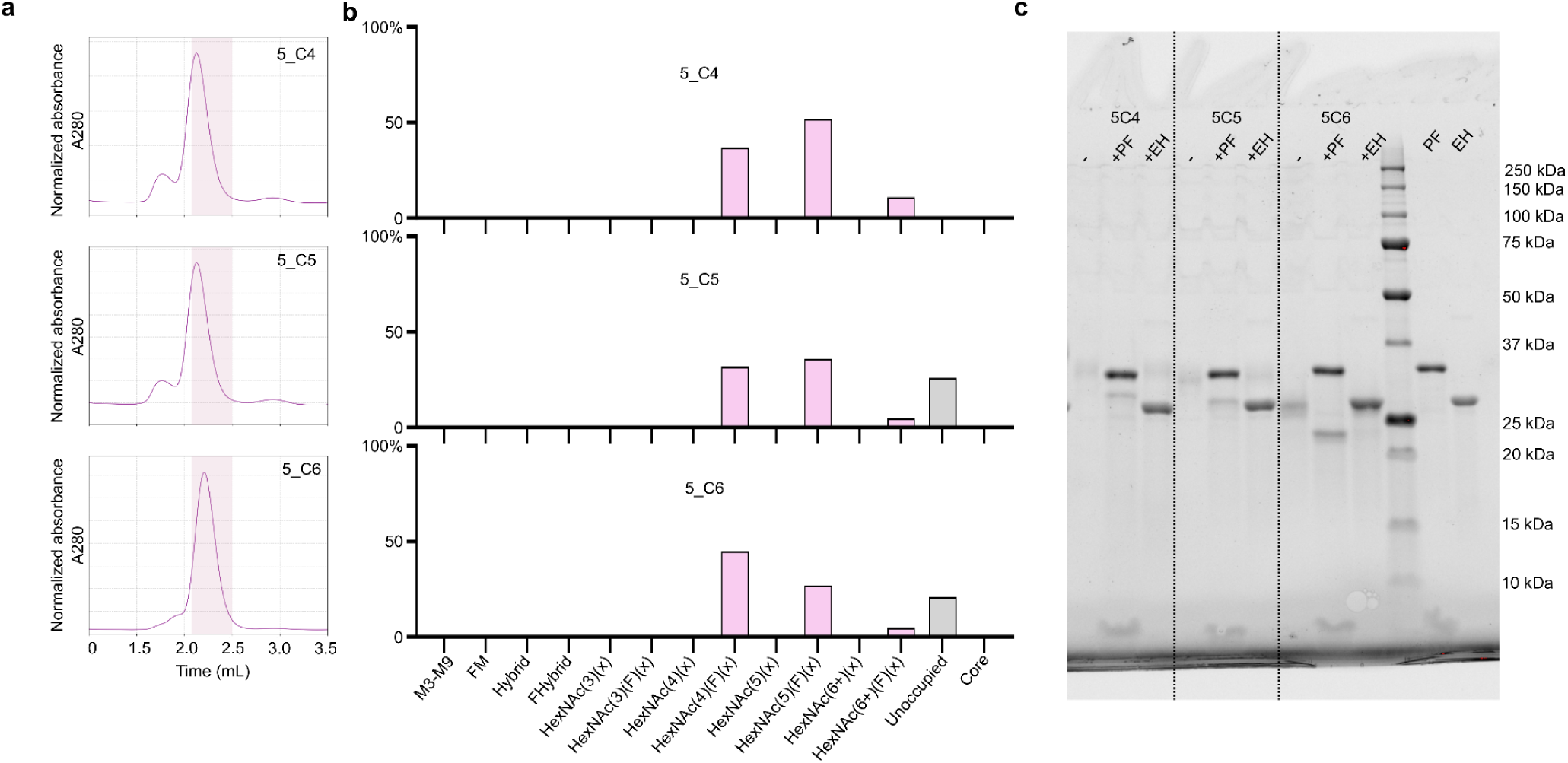
Representative HPLC purification traces and LC-MS/MS extended data for Endo H resistant, PNGase F-sensitive complex glycoproteins 5_C4–C6. a, Normalized HPLC traces from 3 representative *de novo* glycoproteins from top to bottom: 5_C4, 5_C5, and 5_C6. **b**, Compositional site-specific glycan analysis of *de novo* glycoproteins. The graphs summarize quantitative mass spectrometric analysis of the glycan simplified into categories of glycans. The oligomannose-type glycan series (Man9 to Man3) is colored green, afucosylated and fucosylated hybrid-type glycans (hybrid and F hybrid) are dark pink, and complex glycans are grouped according to the number of antennae and presence of core fucosylation and are colored pink. Unoccupancy of an N-linked glycan site is represented in gray. **c,** SDS-PAGE electrophoresis of PNGase F and Endo H digestion. Mobility shift upon treatment with PNGase F suggests presence of glycan, and lack of shift with Endo H suggests complex glycans. Lanes are as follows: **1** Untreated protein 5_C4 **2** Protein 5_C4 + PNGase F **3** Protein 5_C4 + Endo H **4** Untreated Protein 5_C5 **5** Protein 5_C5 + PNGase F **6** Protein 5_C5 + Endo H **7** Untreated protein 5_C6 **8** Protein 5_C6 + PNGase F **9** Protein 5_C6 + Endo H **10** Ladder **11** Pngase F alone **12** Endo H alone.

**Extended Data Figure 2:**
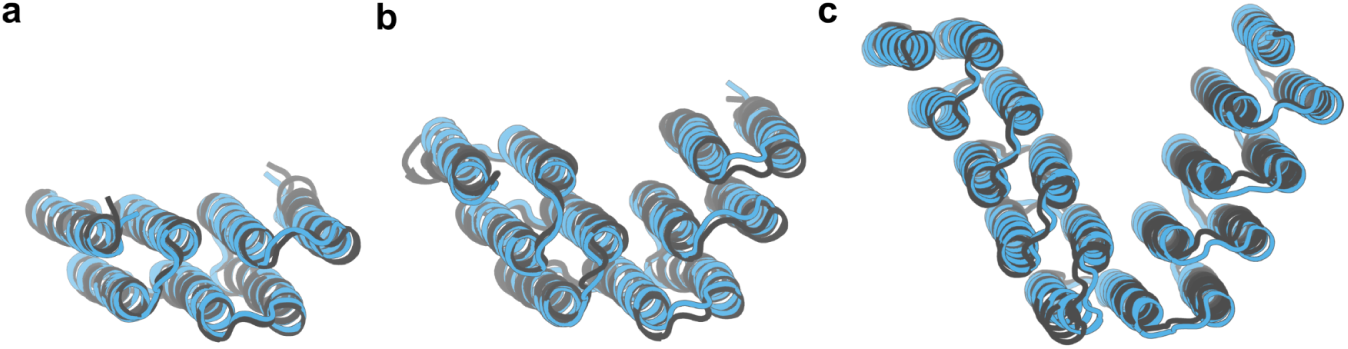
X-ray crystallography determined structures of THR-based proteins. Aligned design structures with resolved crystal structure for THRgly1 **a,** 2 **b**, and 4 **c**. Designs are in blue, solved structures shown in gray.

**Extended Data Figure 3:**
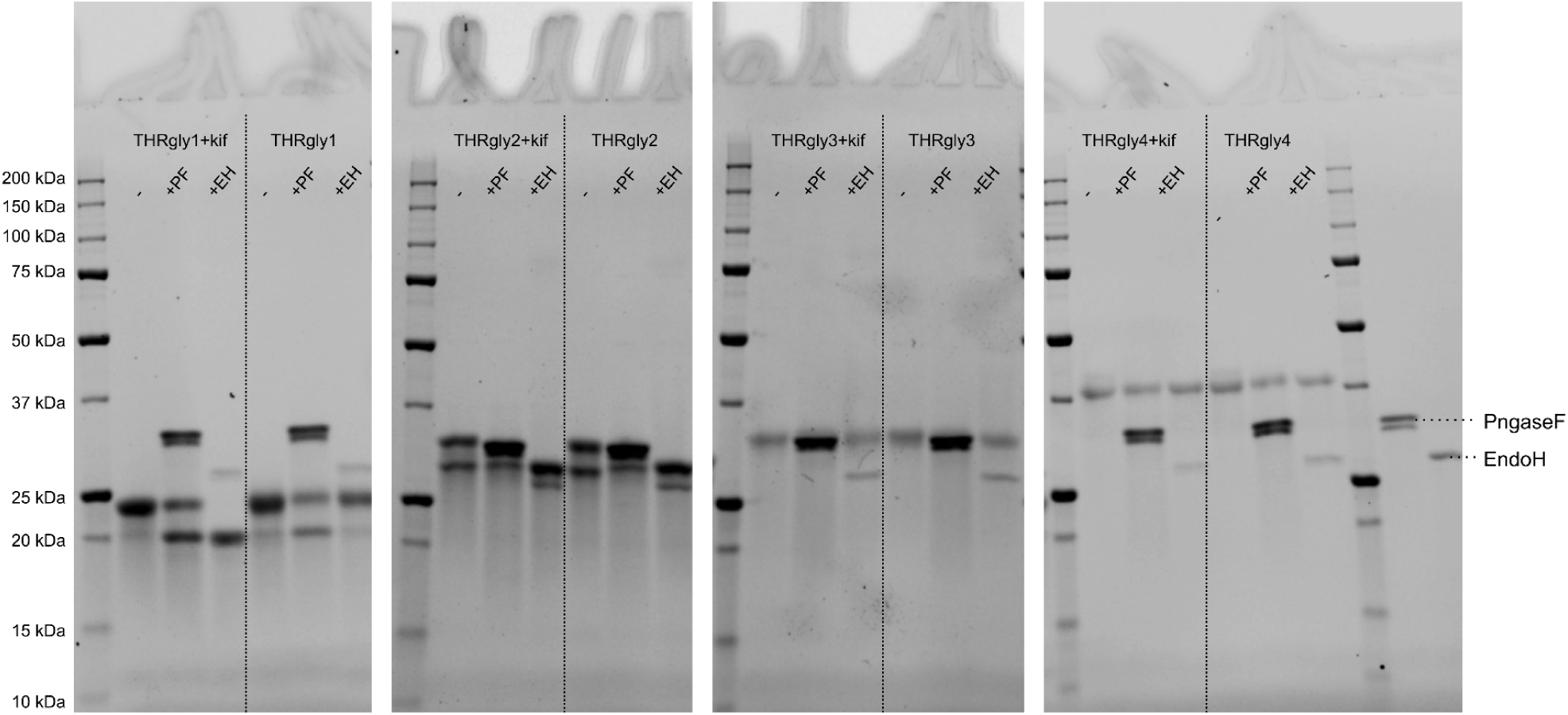
Enzymatic Digestion of THR-based proteins. SDS-PAGE electrophoresis of PNGase F and Endo H digestion. Mobility shift upon treatment with PNGase F suggests presence of glycan, and lack of shift with Endo H suggests complex glycans. Each THRgly variant is shown made in cells with and without kifunensine. Lanes are as follows for each condition: **1**; Untreated protein **2**; Protein + PNGase F **3**; Protein + Endo H. The rightmost two wells show enzymes alone.

**Extended Data Figure 4:**
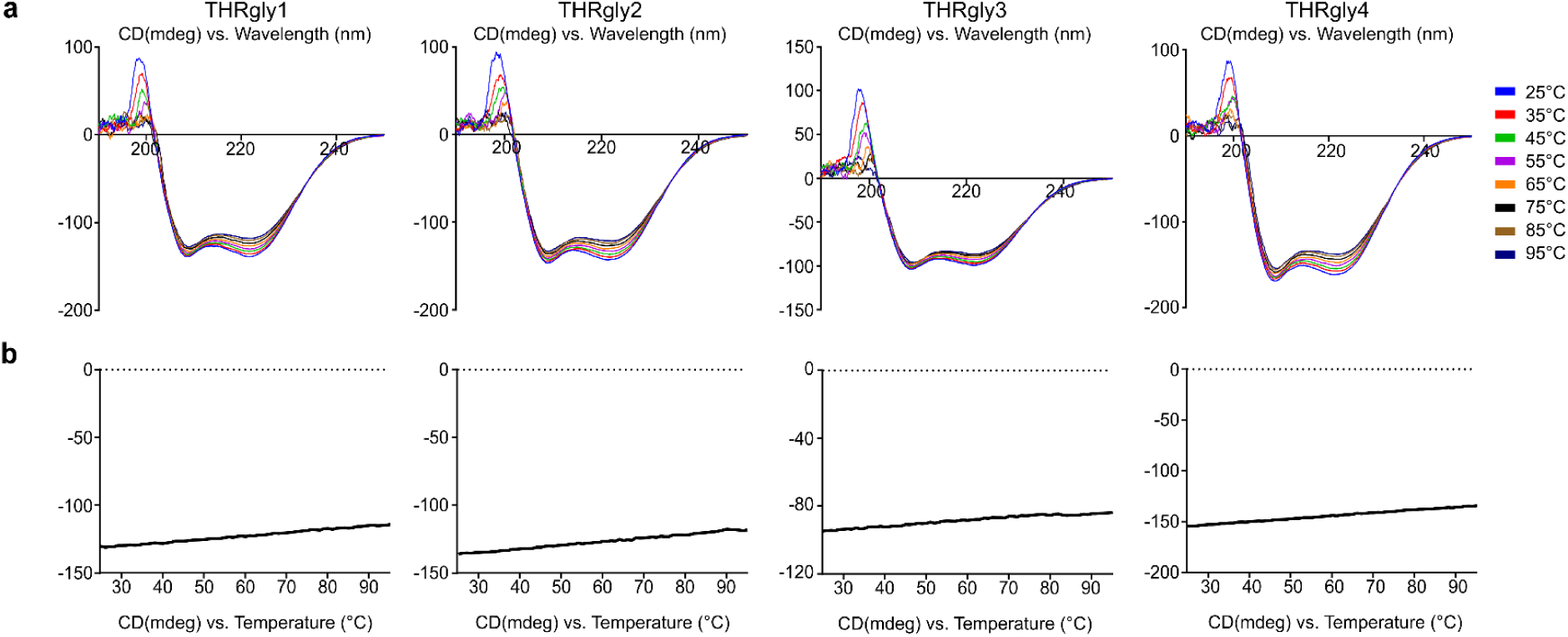
CD Data for THR-based proteins a, Circular dichroism spectrum for THRgly1-4. Full wavelength scan (190nm-250nm) collected every 10°C from 25-95°C. **b,** Thermal denaturation for THRgly1-4 monitored at fixed wavelength (217nm) measurements collected every 1°C from 25-95°C.

**Extended Data Figure 5:**
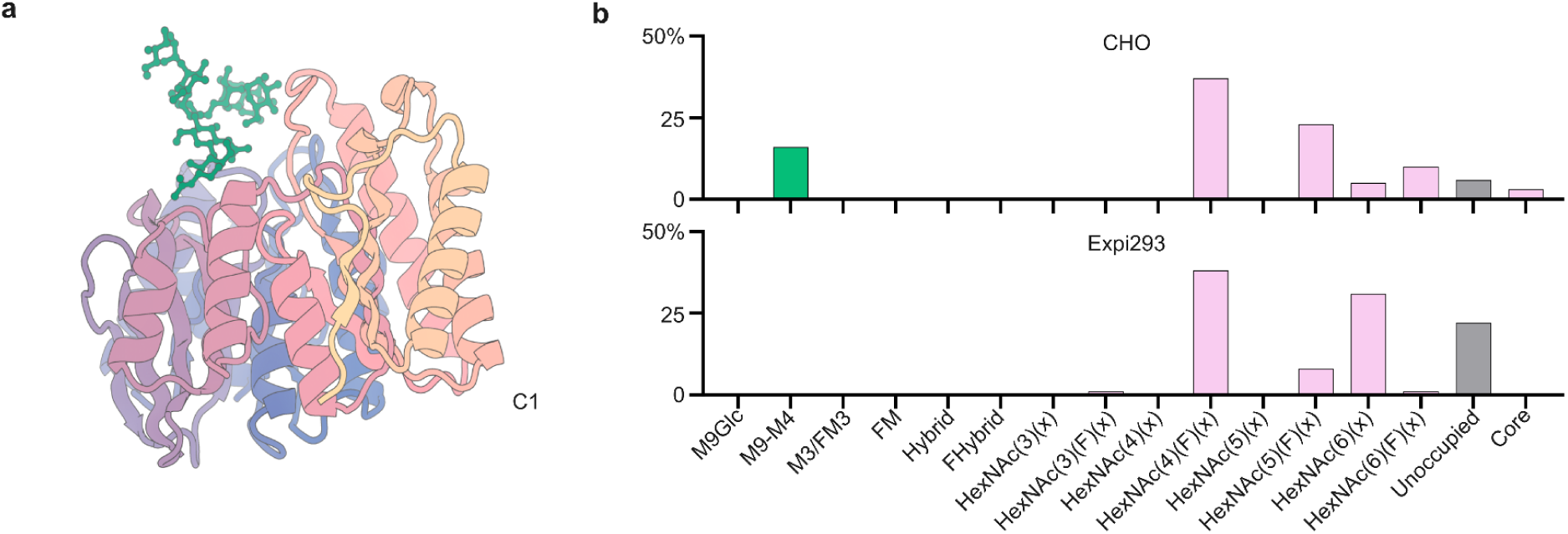
Compositional site-specific glycan analysis of complex control de novo glycoprotein C1 a, AF3 predicted structure modeled for de novo complex glycoprotein C1 used as a control. **b,** Graph displays quantitative mass spectrometric analysis of the glycan simplified into categories of glycans. The oligomannose-type glycan series (Man9 to Man4) is colored green, afucosylated and fucosylated hybrid-type glycans (hybrid and F hybrid) are dark pink, and complex glycans are grouped according to the number of antennae and presence of core fucosylation and are colored pink. Unoccupancy of an N-linked glycan site is represented in gray. Proteins were produced in CHO (top) and Expi293F (bottom) mammalian cells.

**Extended Data Figure 6:**
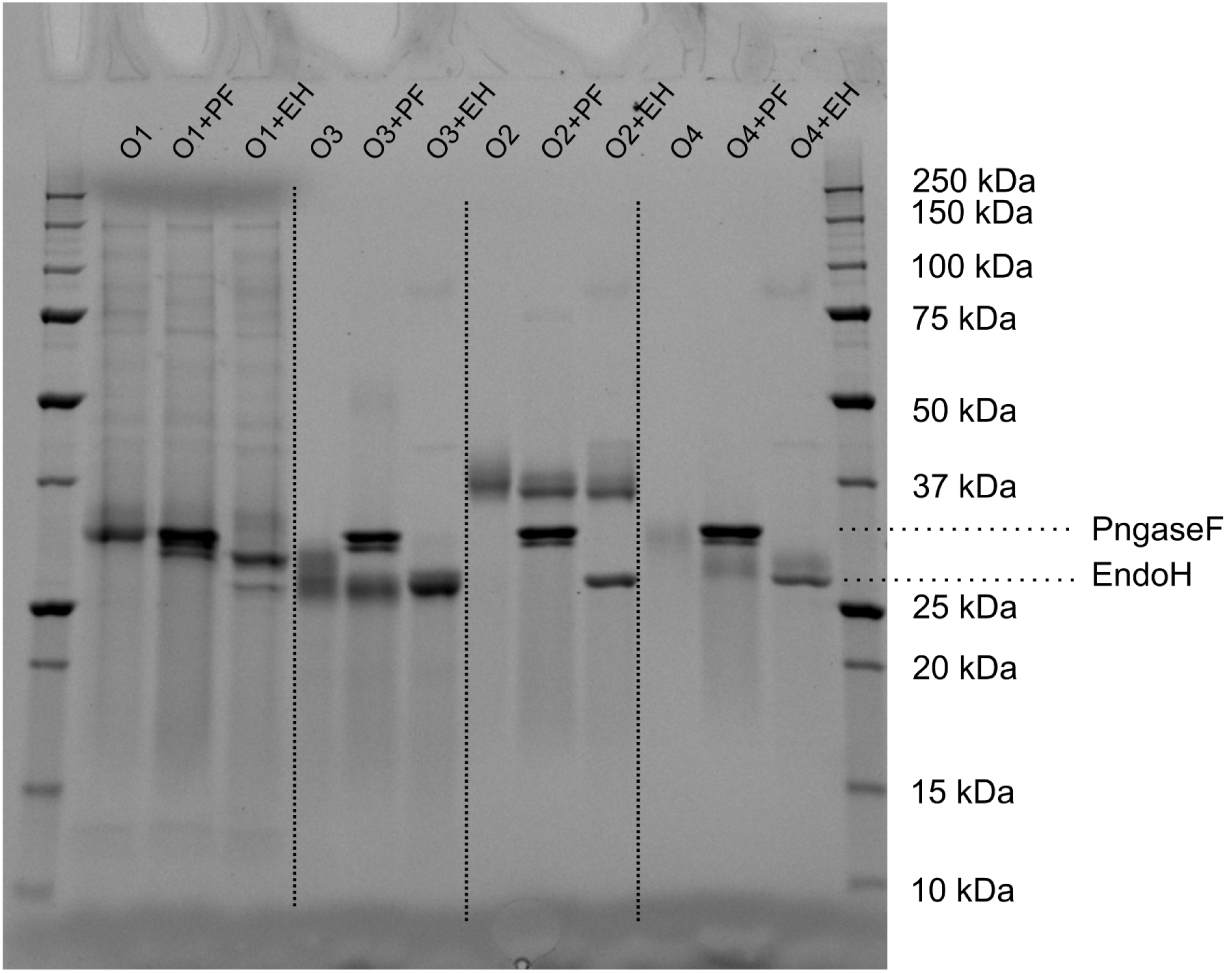
SDS-PAGE gel PNGase F and Endo H digestion for *de novo* oligomannose-type glycoproteins. SDS-PAGE electrophoresis of PNGase F and Endo H digestion. Mobility shift upon treatment with PNGase F suggests presence of glycan, and shift with Endo H suggests oligomannose glycans. Lanes are as follows: **1**; Ladder **2;** Untreated protein O1 **3**; Protein O1 + PNGase F **4**; Protein O1 + Endo H **5**; Untreated Protein O3 **6**; Protein O3 + PNGase F **7**; Protein O3 + Endo H **8**; Untreated protein O2 **9**; Protein O2 + PNGase F **10**; Protein O2 + Endo H **11**; Untreated protein O4 **12**; Protein O4 + PNGase F **13**; ProteinO4 + Endo H **14**; Ladder

**Extended Data Figure 7:**
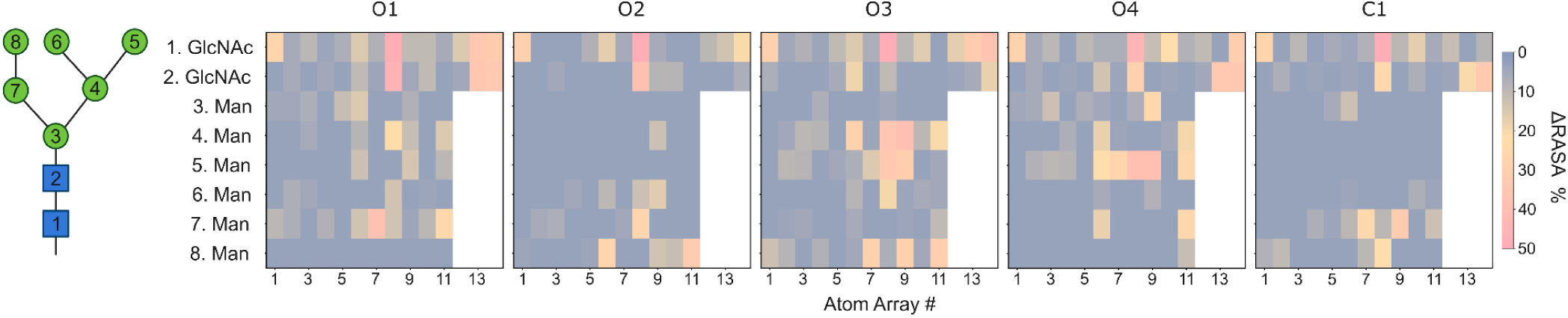
Calculated relative accessible surface area (ΔRASA) scores by atom. ΔRASA scores for four selected oligomannose glycoproteins and one complex glycoprotein. RASA determines relative accessible surface area calculated using a 1.4 Å probe relative to the theoretical maximum accessibility of an isolated atom, 4π(r + r_probe)^2^. Scores are first determined by scoring glycan RASA with protein to determine total RASA, then a second calculation performed with protein atoms removed, capturing self-occlusion by neighboring glycan atoms. ΔRASA is determined by the absolute difference between these values, with higher values indicating greater occlusion by protein structure. Values are shown as percentages because bonded glycan neighbors occlude part of the isolated-atom reference, so ΔRASA cannot approach 100% for a fully occluded atom.

**Extended Data Figure 8:**
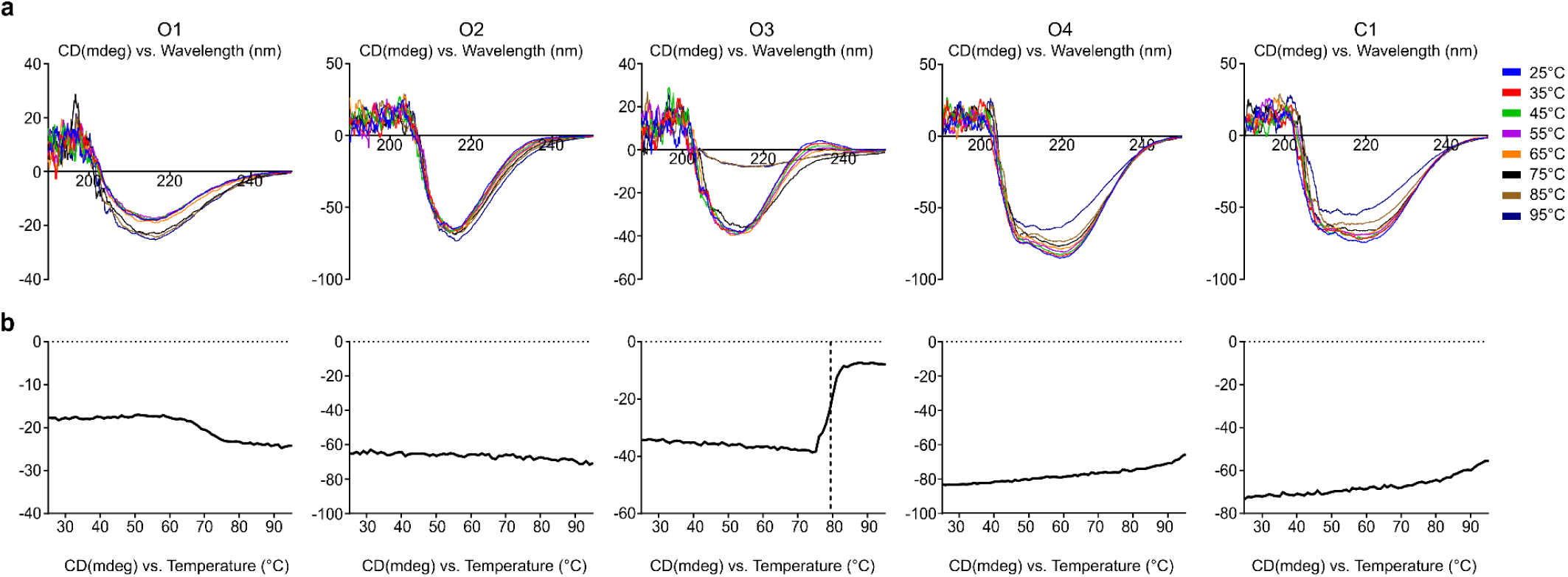
CD data for diffusion-based selected glycoproteins a, Full wavelength scan (190 nm–250 nm) collected every 10°C from 25–95°C (top panels). **b,** Thermal denaturation monitored at fixed wavelength (222 nm or 217 nm, as labeled) measurements collected every 1°C from 25–95°C (bottom panels). O1 monitored at 217 nm showed no clear cooperative unfolding transition up to 95°C. A possible aggregation event between 60–70°C may be indicated by increasing ellipticity. O2 monitored at 217 nm exhibited no clear cooperative unfolding transition up to 95°C. O3 monitored at 217 nm showed a cooperative unfolding transition midpoint temperature of 79°C determined using a four-parameter sigmoidal model in GraphPad Prism. O4 monitored at 222 nm exhibited no clear cooperative unfolding transition up to 95°C. Partial or weak unfolding event possible above 85°C noted by loss of ellipticity with increasing temperature. C1 monitored at 222 nm showed no clear cooperative unfolding transition up to 95°C. Possible partial or weak unfolding event above 80°C noted by loss of ellipticity with increasing temperature.

---

**Extended Data Table 1:** Crystallographic Table.

|  | THRgly1 (PDB ID: 39BF) | THRgly2 (PDB ID: 39BG) | THRgly4 (PDB ID: 39BH) | O4 (PDB ID: 39BI) |
| --- | --- | --- | --- | --- |
| **Data collection** |  |  |  |  |
| Space group | *P* 6<sub>1</sub> 2 2 | *C* 2 2 2<sub>1</sub> | *P* 1 | *P* 2<sub>1</sub> 2<sub>1</sub> 2<sub>1</sub> |
| Cell dimensions |  |  |  |  |
| *a*, *b*, *c* (Å) | 90.28, 90.28, 469.62 | 80.54, 98.78, 153.51 | 47.29, 55.23, 79.88 | 52.48, 109.46, 109.49 |
| $\alpha$, $\beta$, $\gamma$ (°) | 90, 90, 120 | 90, 90, 90 | 100.85, 90.39, 95.70 | 90, 90, 90 |
| Resolution (Å) | 156.54 - 3.57 (3.91 - 3.57) | 76.76 - 3.51 (4.02 - 3.51) | 29.19 - 1.93 (1.98 - 1.93) | 38.70 - 3.90 (4.30 - 3.90) |
| *R*<sub>merge</sub> | 0.794 (1.762) | 0.2593 (2.046) | 0.095 (0.733) | 0.467 (2.208) |
| *I* / σ*I* | 6.9 (2.9) | 5.07 (1.03) | 6.2 (1.0) | 6.2 (1.5) |
| Completeness (%) | 100.0 (100.0) | 99.07 (99.85) | 94.3 (89.7) | 99.8 (99.4) |
| Redundancy | 35.1 (36.3) | 13.2 (12.7) | 2.2 (2.3) | 13.8 (14.0) |
| CC<sub>1/2</sub> | 0.976 (0.497) | 0.994 (0.297) | 0.997 (0.581) | 0.991 (0.706) |
| **Refinement** |  |  |  |  |
| Resolution (Å) | 78.18 - 3.57 (3.7 - 3.57) | 76.76 - 3.51 (4.02 - 3.51) | 29.19 - 1.93 (1.98 - 1.93) | 38.70 - 3.90 (4.30 - 3.90) |
| No. reflections | 14512 (1379) | 7871 (2587) | 55368 (3808) | 6068 (1444) |
| *R*<sub>work</sub> / *R*<sub>free</sub> | 0.2667 (0.3277) / 0.3058 (0.3678) | 0.3295 (0.3525) / 0.3581 (0.3672) | 0.2210 (0.3008) / 0.2681 (0.3414) | 0.2926 (0.3471) / 0.3496 (0.3719) |
| No. atoms |  |  |  |  |
| Protein | 6120 | 4783 | 7412 | 5536 |
| Water | n/a | n/a | 113 | 7 |
| *B*-factors |  |  |  |  |
| Protein | 58 | 55 | 37 | 188 |
| Water | n/a | n/a | 34 | 140 |
| R.m.s. deviations |  |  |  |  |
| Bond lengths (Å) | 0.003 | 0.003 | 0.005 | 0.005 |
| Bond angles (°) | 0.500 | 0.500 | 0.590 | 0.880 |

\*Single Crystal used for each data collection. \*Values in parentheses are for highest-resolution shell.

**Extended Data Table 2:** Prediction Model Comparison Table.

| **Name** | **O1** | **O2** | **O3** | **O4** |
| --- | --- | --- | --- | --- |
| **AF2 Model** | 5 | 2 | 1 | 5 |
| **AF2 Recycles** | 3 | 3 | 3 | 3 |
| **AF2 pLDDT** | 42.7 | 85.6 | 67.7 | 77.5 |
| **AF2 pTM** | 0.3 | 0.8 | 0.7 | 0.8 |
| **AF2 pAE** | 21.5 | 7.5 | 12.6 | 9.2 |
| **AF2 RMSD to design** | 21.8 | 0.7 | 16.8 | 6.1 |
| **RF3 pLDDT** | 87.2 | 82.2 | 81.2 | 75.4 |
| **RF3 pAE** | 6.5 | 8.8 | 8.8 | 11.4 |
| **RF3 RMSD to design** | 1.50 | 2.82 | 2.26 | 7.60 |
| **AF3 Model** | 1 | 0 | 0 | 0 |
| **AF3 pLDDT** | 84.0 | 81.9 | 82.8 | 75.4 |
| **AF3 pAE** | 7.8 | 9.3 | 8.6 | 10.3 |
| **AF3 pTM** | 0.85 | 0.83 | 0.83 | 0.78 |
| **AF3 RMSD to design** | 1.2 | 3.1 | 2.1 | 5.7 |

